# Mitochondrial genomes reveal maternal lineages of Late Iron Age sheep (*Ovis aries*) in Denmark

**DOI:** 10.64898/2026.04.19.719464

**Authors:** Jonas Holm Jæger, Valeria Mattiangeli, Jens Ulriksen, Torben Sarauw, Mads Dengsø Jessen

## Abstract

Sheep husbandry played an important role in the agrarian and textile economies of Late Iron Age Denmark, yet the genetic structure of sheep populations from this period remains poorly understood. In this study, we analyse complete mitochondrial genomes from five Late Iron Age sheep recovered from four Danish archaeological sites dating to the fifth to tenth centuries AD.

All sequenced sheep belong to mitochondrial haplogroup B and fall within the B1a lineage, the predominant maternal lineage in European sheep. Each individual represents a distinct haplotype, resulting in high haplotype diversity despite the limited sample size. Population differentiation between Danish ancient sheep and modern reference populations from Fennoscandia and northwest Europe is very low, indicating limited maternal genetic differentiation at the regional scale. Median-joining network analysis further shows that Danish haplotypes are distributed across the broader northern European haplogroup B lineage rather than forming a geographically distinct cluster.

These results suggest that Danish sheep populations during the Late Iron Age maintained multiple maternal lineages and were embedded within wider northern European genetic networks. The observed mitochondrial diversity is consistent with sheep husbandry systems that were not restricted to narrow maternal breeding stocks during a period associated with expanding textile production.

**Highlights:**

- Complete mitogenomes from Danish Late Iron Age sheep
- High maternal diversity within haplogroup B
- Genetic links between Late Iron Age and modern sheep

## 1 Introduction

Sheep (*Ovis aries*) were domesticated in Southwest Asia by approximately 8,000 BC (Daly et al., 2025; Kaptan et al., 2024; Vigne, 2011; Zeder, 2008) and subsequently dispersed across Europe through a series of migratory and cultural processes. By around 4,000 BC, domesticated sheep had reached Southern Scandinavia, where early husbandry practices appear to have focused primarily on producing meat and dairy products (Groß et al., 2024; Sørensen, 2014). From their initial introduction, Scandinavian sheep populations were shaped by human mobility, environmental constraints, and shifting economic demands.

One of the most significant transformations occurred during the second millennium BC, when novel, “improved” woolly sheep were introduced into northern Europe, ultimately replacing or admixing with earlier, more “primitive” Neolithic populations and facilitating the development of specialised textile economies (Chessa et al., 2009; Rannamäe et al., 2020; Schroeder et al., 2017). During the Late Iron Age, spanning the Germanic Iron Age (AD 375/400-750) and Viking Age (AD 750-1050), archaeological evidence indicates an intensification of wool-based textile production, reflected in increasing numbers of textile tools and the emergence of large settlement complexes with numerous pithouses associated with textile production (Andersson Strand, 2021).

Despite extensive palaeogenetic research on the domestication and dispersal of sheep across Eurasia (Daly et al., 2025; Deng et al., 2020; Kaptan et al., 2024; Morell Miranda et al., 2023; Yurtman et al., 2021), Scandinavia remains markedly underrepresented in ancient DNA research. Denmark, in particular, lacks genomic coverage for later prehistoric periods. To date, only two ancient sheep mitogenomes have been reported from Denmark: one from an Early Neolithic individual from Havnø in northern Jutland (3708-3526 cal. BC) and one from a Late Roman Iron Age (AD 250/260-310/320) individual from Storskoven on Zealand (Brandt, 2014). Consequently, maternal lineage diversity and connectivity among Danish sheep during the Late Iron Age remain poorly understood.

In this study, ancient sheep mitogenomes from Denmark, dating to the Germanic Iron Age and Viking Age, are analysed to evaluate patterns of maternal lineage diversity during a period of substantial economic and social transformation. The analysis examines whether shifts in sheep exploitation and textile craft specialisation documented in the archaeological record were accompanied by detectable changes in mitochondrial haplogroup composition and patterns of maternal lineage diversity. By comparing newly generated mitogenomes with published ancient and modern sheep mitochondrial datasets, this study provides the first systematic assessment of mitochondrial diversity of Danish sheep during the Late Iron Age. In doing so, it establishes a genetic baseline for later prehistoric sheep husbandry in Southern Scandinavia and contributes to broader discussions of livestock mobility, breeding practices, and economic organisation in northern Europe.

## 2 Materials and methods

### 2.1 Archaeological samples

A total of six ancient sheep petrous bones were selected from four Germanic Iron Age and Viking Age archaeological sites in Denmark: Vester Egesborg (Ulriksen, 2018), Fugledegård (Jørgensen et al., 2019), Bejsebakken (Sarauw, 2019), and Aggersborg (Roesdahl et al., 2014) (Figure 1; Table 1; Supplementary S1). These sites represent a range of settlement types, including a landing and assembly site (Vester Egesborg), an elite residential complex (Fugledegård), a Viking Age ring fortress (Aggersborg), and a Late Iron Age specialised craft and trading site (Bejsebakken). Bejsebakken and Aggersborg are located in the Limfjord region of northern Jutland, while Fugledegård and Vester Egesborg are situated in western and southern Zealand, respectively. The selected specimens represent geographically distributed sheep remains from northern and eastern Denmark. All specimens were morphologically identified as sheep or ovicaprine before analysis and exhibited good macroscopic preservation.

**Table 1.** The archaeological context of the six ancient sheep specimens included in this study. The samples derive from Late Iron Age contexts within present-day Denmark.

| ID | Site | Morphology | Context | Date | Archaeological period | Reference |
| --- | --- | --- | --- | --- | --- | --- |
| JJ1 | Vester Egesborg | *Ovis/Capra* | Pit-house | AD 6-10th cent. | Late Germanic Iron Age-Viking Age | (Enghoff, 2018) |
| JJ2 | Fugledegård | *Ovis aries* | Royal residence | AD 8-10th cent. | Late Germanic Iron Age-Viking Age | Anne Birgitte Gotfredsen, unpublished |
| JJ3 | Bejsebakken | *Ovis/Capra* | Pit | AD 5-6th cent. | Late Germanic Iron Age | (Kveiborg, 2019) |
| JJ4 | Bejsebakken | *Ovis/Capra* | Pit-house | AD 7-10th cent. | Late Germanic Iron-Viking Age | (Kveiborg, 2019) |
| JJ5 | Aggersborg | *Ovis/Capra* | Pit-house | AD 9-10th cent. | Viking Age | (Rosenlund & Hatting, 2014) |
| JJ6 | Aggersborg | *Ovis/Capra* | Pit-house | AD 9-10th cent. | Viking Age | (Rosenlund & Hatting, 2014) |

**Figure 1.**
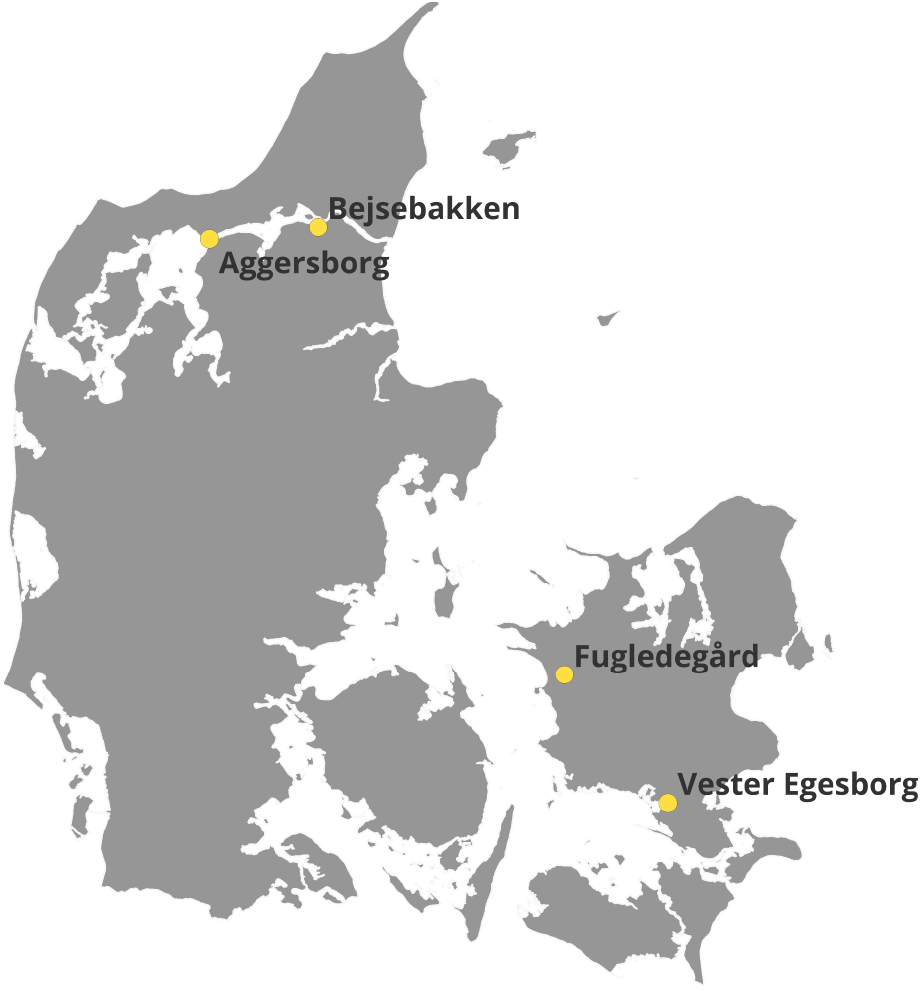
Map of archaeological sites. Geographic distribution of Late Iron Age archaeological sites included in this study: Vester Egesborg, Fugledegård, Bejsebakken, and Aggersborg.

### 2.2 Ancient DNA extraction and library preparation

Petrous bones morphologically identified as sheep were processed in dedicated ancient DNA clean lab facilities at the GLOBE Institute, University of Copenhagen, Denmark, and the Smurfit Institute of Genetics, Trinity College Dublin, Ireland.

#### 2.2.1 Sampling

Subsampling of petrous bones was performed at dedicated ancient DNA facilities at the GLOBE Institute, University of Copenhagen, Denmark. Subsampling was performed using a Dremel Multitool equipped with a diamond split disc. Tools and work surfaces were cleaned between samples following established best-practice guidelines for ancient DNA (Pääbo et al., 2004). Subsamples were subsequently powdered using a mixer mill, and 113-126 mg of bone powder was used for downstream DNA extraction.

#### 2.2.2 DNA extraction

DNA extraction was carried out following the protocol described by Gamba et al. (2014), based on (MacHugh et al., 2000; Yang et al., 1998) with minor modifications.

Bone powder was treated with a 0.5% sodium hypochlorite solution (Boessenkool et al., 2016), briefly vortexed, and rotated for 15 minutes. Samples were then washed with 1 mL molecular-grade water and centrifuged at 13,300 rpm prior to removal of the supernatant; this wash step was repeated twice.

Following bleach treatment, samples were washed with EDTA by adding 990 μL of 0.5 M EDTA, vortexing, and incubating for 30 minutes at 37 °C. After centrifugation at 13,300 rpm, the EDTA was discarded. The remaining pellet was incubated for 12 hours at 37 °C in 990 μL extraction buffer (0.47 M EDTA, 1.7% N-lauroylsarcosine, 0.02 M Tris-HCl, 0.65 U proteinase K) with gentle agitation using an Eppendorf ThermoMixer.

Following incubation, samples were centrifuged at 13,300 rpm for 10 minutes. The supernatant was transferred to Falcon tubes containing 13 mL binding buffer (PB) (Dabney et al., 2013) and processed using Roche reservoir columns. Columns were centrifuged for 4 minutes at 800 x g, rotated 90°, and centrifuged for an additional 2 minutes at 800 x g.

DNA purification was performed using Roche reservoir columns, following the manufacturer’s protocol. DNA was then eluted in a 40 μL EB buffer supplemented with Tween-20 to a final concentration of 0.05%. Extraction blanks were processed alongside samples and sequenced to monitor potential contamination.

#### 2.2.3 UDG treatment

For uracil-DNA glycosylase treatment, 16.25 μL of purified DNA was incubated with 5 μL USER enzyme (1 U/μL; New England BioLabs) for 1 hour at 37 °C, followed by blunt-end repair according to Meyer and Kircher (2010).

#### 2.2.4 Library preparation

Double-stranded DNA libraries were prepared using 21.25 μL USER-treated DNA following Meyer and Kircher (2010), with minor modifications: SPRI purification was replaced with Qiagen MinElute purification, and a heat-inactivation step for Bst polymerase (20 minutes at 80 °C) was introduced, replacing an additional purification step described by Gamba et al. (2014).

PCR indexing was carried out on a total volume of 25 μL containing 20.5 μL of AccuPrime Pfx Polymerase (Invitrogen), 0.5 μL of 10 μM P5 primer, 1 μL of 5 μM P7 primer, and 3 μL of DNA library.

Amplification was performed in a physically separated post-PCR facility using the following cycling conditions: 95 °C for 5 minutes; 12 cycles of 95 °C for 15 seconds, 60 °C for 30 seconds, and 68 °C for 30 seconds; followed by a final extension at 68 °C for 5 minutes. PCR products were purified using Qiagen MinElute spin columns and eluted in a 12 μL EB buffer. Library and PCR blanks were included throughout for contamination control.

#### 2.2.5 Screening of ancient DNA libraries

Ancient DNA libraries and controls were quantified using an Agilent TapeStation 2200 with the D1000 assay kit.

Endogenous DNA content was assessed through low-depth sequencing of each library and control on an Illumina NovaSeq 6000 platform (paired-end, 2×100 bp) at the Genomics Core Facility (TrinSeq), Trinity College Dublin, Ireland.

## 3 Bioinformatics

### 3.1 Archaeological samples (JJ1-JJ6)

Adapters were removed from raw sequencing reads using AdapterRemoval v2.3.3 (Schubert et al., 2016) with the following parameters: --minlength 30 --trimns --trimqualities. Read quality and fragment length distributions were assessed before and after trimming using FastQC (Andrews 2010) to confirm successful adapter removal. Fastq Screen v. 0.14.0 (Wingett & Andrews, 2018) was used to confirm all specimens as sheep using a cohort of reference animal mitogenomes (cattle, sheep, goat, deer, human, pig, dog, horse).

Following trimming, reads were mapped to the *Ovis aries* nuclear reference genome (ARS-UI_Ramb_v2.0; GCA_016772045.1) downloaded from NCBI. Mapping was carried out using bwa aln (Li & Durbin, 2009) with parameters optimised for ancient DNA (-l 1024 -n 0.01 -o 2). The resulting .sai files were converted to SAM format using bwa samse and subsequently to BAM format using SAMtools v1.21 (Danecek et al., 2021). BAM files were filtered to remove unmapped reads and those with mapping quality below 25, then sorted by coordinate.

Duplicate reads were removed using Picard v3.4.0 with the parameters OPTICAL_DUPLICATE_PIXEL_DISTANCE=2500 REMOVE_DUPLICATES=true. Read groups were added to each BAM file using Picard.

For mitochondrial analyses, trimmed reads were aligned to a circularised *Ovis aries* mitochondrial reference genome (NC_001941.1) downloaded from NCBI. Circularisation was done by appending the first 30 bp of the reference sequence to its end to mitigate edge effects during alignment. Alignment, filtering, sorting, and duplicate removal were performed using the same procedure as described above.

### 3.2 Reference samples

Reference datasets were obtained from the European Nucleotide Archive (ENA) and GenBank and included paired-end and single-end sequencing reads, as well as pre-assembled FASTA sequences. Reference samples available only as mitochondrial FASTA sequences were aligned directly to the circularised mitochondrial reference genome before downstream analyses.

Ancient reference reads were aligned to the circularised mitochondrial reference genome using bwa aln, while modern reads were aligned using bwa mem (Li & Durbin, 2009). Post-alignment processing, including sorting, filtering, and duplicate removal, was performed using SAMtools and Picard, following the same procedures described above.

### 3.3 DNA damage and authenticity

Post-mortem DNA damage patterns were assessed for all newly generated Late Iron Age/Viking Age samples using mapDamage2/2.2.2 (Jónsson et al., 2013). Analyses were performed on deduplicated BAM files aligned to the *Ovis aries* nuclear reference genome to characterise fragment length distributions and nucleotide misincorporation patterns.

As libraries were prepared using full UDG treatment, deamination-derived misincorporations were expected to be substantially reduced relative to untreated ancient DNA. Authentication was therefore based on residual enrichment of terminal C→T substitutions at 5′ read ends and complementary G→A substitutions at 3′ ends, together with short fragment length distributions.

### 3.4 Biological sex

Biological sex was inferred from relative X-chromosome coverage using *karyotype.R* (https://github.com/teasdalm/karyotypeR), which implements the approach described by Skoglund et al. (2015). Per-chromosome mapped read counts were obtained from deduplicated BAM files using *samtools idxstats* and normalised by chromosome length. Biological sex was then assigned based on the ratio of X chromosome to autosomal coverage.

Individuals with X-to-autosome ratios approximating ~1.0 were classified as biologically female (XX). In contrast, ratios near 0.5 were classified as male (XY), reflecting differences in X-chromosome copy number between males and females. Diagnostic coverage plots were also generated for each individual to assess classification robustness (Supplementary S2).

### 3.5 Determining mitochondrial haplogroups

Mitochondrial haplogroups were determined for the newly generated Late Iron Age/Viking Age samples. Following mapping to the circularised *Ovis aries* mitochondrial reference genome and standard post-mapping processing, consensus mitochondrial sequences were generated from deduplicated BAM files using ANGSD v.0.9.40 (Korneliussen et al., 2014) and exported in FASTA format. Haplogroup assignment was then performed using MitoToolPy (Peng et al., 2015) with the parameters: *-s sheep -r whole*.

For previously published archaeological and modern reference samples included in the comparative dataset, mitochondrial haplogroups were retained as reported in the original publications. Where necessary, haplogroup nomenclature was harmonised to ensure consistency across datasets.

### 3.6 Nucleotide diversity, neutrality statistics and population differentiation

Nucleotide diversity statistics were calculated using DnaSP v. 6.12.03 (Rozas et al., 2017). Analyses were restricted to sites that were callable in all individuals to ensure comparability across sequences. Following exclusion of sites containing gaps or missing data, a total of 16,480 mitochondrial positions were retained for downstream analyses.

Standard summary statistics, including the number of segregating sites (*S*), number of haplotypes (*h*), haplotype diversity (*Hd*), nucleotide diversity (*π*), and the average number of pairwise nucleotide differences (*k*), were calculated. Neutrality tests were performed using Tajima’s D and Fu’s Fs statistics. Tajima’s D was calculated based on the total number of segregating sites, while Fu’s Fs was calculated using the total number of mutations. Statistical significance was assessed using the default parameters implemented in DnaSP.

Population differentiation (*F*_*ST*_) between Danish ancient sheep and reference populations was estimated in DnaSP using the Hudson, Slatkin and Maddison (1992) estimator for haploid sequence data. Analyses were performed on the same gap-filtered alignment used for diversity statistics. Comparative reference sets were defined as modern Fennoscandia (Norway, Sweden, and Finland) and modern Northwest Europe (the Netherlands, Germany, Britain, and associated North Sea-region breeds). Significance was assessed using permutation tests implemented in DnaSP with 1,000 replicates. Negative *F*_*ST*_ estimates were interpreted as zero differentiation. Because mitochondrial genomes are nonrecombining and maternally inherited, differentiation statistics are reported as measures of maternal population structure, and gene flow parameters derived from *F*_*ST*_ (e.g. *Nm*) are not interpreted due to their potential bias and high variance under conditions of low differentiation (Hudson et al., 1992).

### 3.7 Median-joining networks

Complete mitochondrial genome consensus sequences were aligned using MAFFT v7.526 (Katoh & Standley, 2013). The resulting alignment formed the basis for all downstream diversity analyses and network construction.

Relationships between mitochondrial haplotypes were visualised in a median-joining network implemented in PopART (Leigh & Bryant, 2015). Complete mitochondrial genome sequences included in the network were compiled from this study and previously published datasets. A full list of sequences is provided in Supplementary S3.

Gaps and missing data were treated as missing characters. Nodes represent unique haplotypes, with node size scaled to haplotype frequency, and hatch marks indicating the number of mutational steps separating haplotypes. Inferred median vectors (unsampled or extinct intermediate haplotypes) are shown as small intermediate nodes.

## 4 Results

### 4.1 DNA preservation, coverage, and authenticity

A total of six ancient sheep samples were processed for mitochondrial genome analysis. Of these, five yielded sufficient endogenous DNA content and adequate coverage for downstream analyses. At the same time, one sample (JJ5 from Aggersborg) failed to produce usable mitochondrial data and was excluded from further analyses. For the five retained samples, endogenous DNA proportions ranged from 13.9% to 63.5%, with mean read lengths of 49-54 bp, matching expectations for ancient DNA (Supplementary S4).

Post-mortem DNA damage patterns were assessed using mapDamage2/2.2.2 (Jónsson et al., 2013). In line with full UDG treatment, terminal misincorporation rates were reduced relative to untreated ancient DNA; however, all samples exhibited residual enrichment of 5’ C→T and complementary 3’ G→A substitutions at read termini, together with short fragment length distributions characteristic of ancient DNA. These patterns support the authenticity of the sequenced samples (Supplementary Figure S4).

After removing duplicates, the number of mapped reads varied substantially among individuals, resulting in mitochondrial genome coverage ranging from 36.6× to 65.0×. Nuclear genome coverage was low across all samples (0.07-0.27×). Both male and female individuals were represented among the successfully sequenced samples.

Summary statistics for sequencing, coverage, and haplogroup assignment for all analysed samples are provided in Table 2:

**Table 2.** Sequencing and mapping summary statistics for ancient sheep samples included in this study. Mitochondrial haplogroups are based on reconstructed mitochondrial genomes. Sample JJ5 was excluded from downstream analyses due to negligible endogenous DNA content and insufficient mitochondrial coverage.

| ID | site | raw reads | trimmed reads | aligned > Q25 reads | duplicates removed | endogenous % | average read length | bases mapped | nuclear coverage (x) | mitochondrial coverage (x) | sex | mitochondrial haplogroup |
| --- | --- | --- | --- | --- | --- | --- | --- | --- | --- | --- | --- | --- |
| JJ1 | Vester Egesborg | 22952775 | 20904657 | 13265884 | 1411045 | 63.46 | 53 | 706066164 | 0.27 | 65.02 | M | B1a1 |
| JJ2 | Fugledegård | 26271148 | 23249683 | 13635777 | 1820888 | 58.65 | 52 | 702511021 | 0.27 | 43.46 | F | B1a1b, B1a2a1 |
| JJ3 | Bejsebakken | 27846570 | 24805718 | 3436403 | 9424612 | 13.85 | 54 | 185807207 | 0.07 | 36.60 | F | B1a1b |
| JJ4 | Bejsebakken | 32782077 | 28110528 | 11050331 | 954475 | 39.31 | 52 | 570879413 | 0.22 | 45.03 | F | B1a1b, B1a2a1 |
| JJ5 | Aggersborg | 19930330 | 15383393 | 76955 | 86545 | 0.50 | 55 | 4238362 | 0.00 | 0.42 | M | - |
| JJ6 | Aggersborg | 27727796 | 23893767 | 10966831 | 1532117 | 45.90 | 49 | 540527201 | 0.21 | 43.71 | M | B1a1b |

### 4.2 Mitogenome recovery and haplogroup composition

Complete mitochondrial genomes were successfully reconstructed for five of the six analysed samples. All successfully sequenced individuals were assigned to mitochondrial haplogroup B, with sub-haplogroups B1a1 and B1a1b predominating. Two individuals (JJ2 and JJ4) displayed ambiguity at diagnostic positions and could be assigned to either B1a1b or the derived sub-haplogroup B1a2a1, likely reflecting within-lineage variation or limited coverage at diagnostic positions.

Haplogroup assignments were consistent across sites and sexes, with no clear association between mitochondrial lineage and geographic location within the dataset. Multiple individuals shared closely related haplotypes within haplogroup B1a1b, including samples from different sites, indicating overlapping maternal lineages across the study area. No mitochondrial haplogroups outside haplogroup B were observed among the analysed samples.

### 4.3 Mitochondrial diversity

Mitochondrial diversity among Danish Late Iron Age sheep was assessed using standard summary statistics based on five mitochondrial genomes (Table 3). All five individuals possessed distinct haplotypes (*h* = 5), yielding a haplotype diversity of 1.00 ± 0.126. A total of 47 segregating sites were identified across the dataset, with a nucleotide diversity (*π*) of 0.00123 and an average of 20.2 pairwise differences between sequences. Neutrality tests yielded a negative Tajima’s D (*D* = −0.789, *p* > 0.10) and a positive Fu’s Fs value (*Fs* = 0.533, *p* > 0.10), with neither statistic differing significantly from expectations under neutrality. Mitochondrial diversity statistics were calculated across 16,480 callable mitochondrial positions shared among all Danish Late Iron Age samples.

**Table 3.** Mitochondrial diversity statistics for Danish Late Iron Age sheep. Summary statistics were calculated from five complete mitochondrial genomes using 16,480 callable positions shared across all individuals. Reported metrics include the number of callable sites (bp), the number of segregating sites (S), haplotype count (h), haplotype diversity (Hd), nucleotide diversity (π), average pairwise differences (k), and neutrality statistics (Tajima’s D and Fu’s Fs). Neutrality tests are reported for completeness and should be interpreted cautiously, given the small sample size.

| | *n* | *bp* | *S* | *h* | *Hd* ± SE | $\pi$ | *k* | Tajima's D | Fu's $F_s$ |
| --- | --- | --- | --- | --- | --- | --- | --- | --- | --- |
| This study | 5 | 16,480 | 47 | 5 | $1.00 \pm 0.126$ | 0.00123 | 20.2 | -0.789 ($p > 0.10$) | 0.533 ($p > 0.10$) |

### 4.4 Population differentiation (*F*_*ST*_)

Pairwise mitochondrial differentiation between populations was low to near-zero. *F*_*ST*_ estimates between Danish ancient sheep and modern Fennoscandian reference populations were 0.004, and between Danish ancient sheep and modern Northwest European reference populations were approximately zero. *F*_*ST*_ between modern Fennoscandia and Northwest Europe was 0.020 (see Table 4; Supplementary S5). Although differentiation is numerically lower between Danish and Northwest European samples, this difference is minimal and not interpreted further.

**Table 4.** Pairwise mitochondrial population differentiation. Pairwise F_ST_ estimates calculated using complete mitochondrial genomes. Negative F_ST_ values were interpreted as zero differentiation.

| POPULATION 1 | POPULATION 2 | $F_{ST}$ |
| --- | --- | --- |
| Ancient Danish sheep | Modern Fennoscandia | 0.004 |
| Ancient Danish sheep | Modern Northwest Europe | ~ 0 |
| Modern Fennoscandia | Modern Northwest Europe | 0.020 |

### 4.5 Median-joining network analysis

Relationships among complete mitochondrial genomes were visualised using a median-joining network constructed from the aligned dataset of ancient and modern sheep (Figure 2). Median-joining networks are widely used to explore intraspecific mitochondrial variation and to visualise relationships among closely related haplotypes (Bandelt et al., 1999). The network is dominated by haplogroup B, characterised by an interwoven topology with short mutational distances among haplotypes compatible with patterns observed in ancient Baltic sheep populations (Rannamäe et al. 2016; Rannamäe et al. 2016).

**Figure 2.**
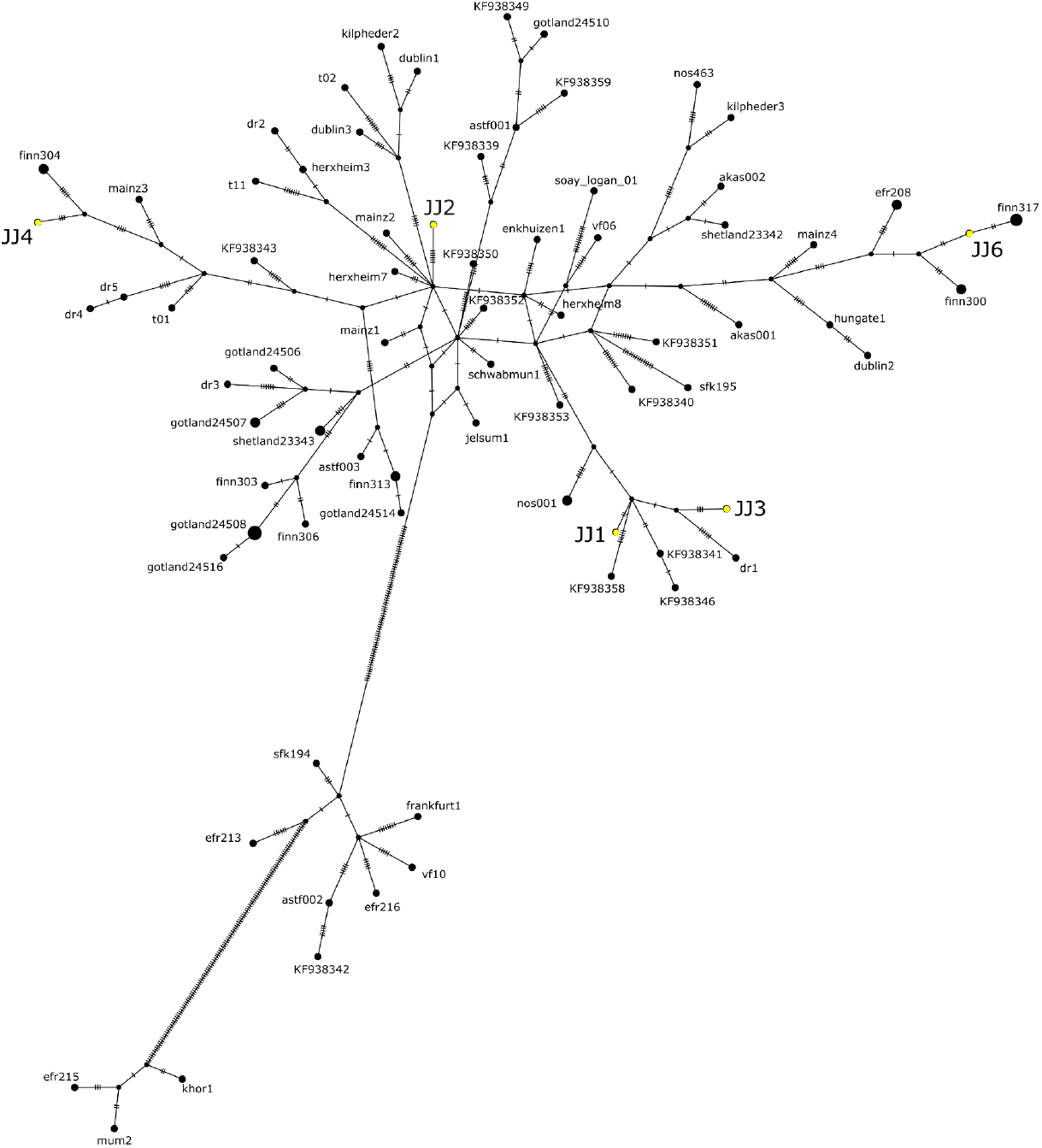
Median-joining network of mitochondrial genomes. Median-joining network showing relationships among complete mitochondrial genomes of ancient and modern sheep included in this study. Danish Late Iron Age samples are highlighted in yellow. Each circle represents a unique haplotype, with circle size proportional to haplotype frequency. Lines connecting nodes represent mutational steps, with hatch marks indicating the number of nucleotide differences between haplotypes. Small black nodes represent inferred median vectors corresponding to unsampled or extinct haplotypes. Danish Late Iron Age samples are distributed within the broader haplogroup B1a component alongside Scandinavian, Northwest European, and Eurasian reference haplotypes.

All Danish mitogenomes fall within the haplogroup B1a subgroup and are dispersed throughout the broader northern European B lineage rather than forming a geographically discrete Scandinavian cluster. Haplogroup B has previously been identified as the predominant maternal lineage in European sheep populations (Hiendleder et al., 1998; Meadows et al., 2007). Samples JJ1 (Vester Egesborg), JJ3 (Bejsebakken) and JJ6 (Aggersborg) occur within the principal B1a component and connect via short paths through inferred median vectors to Scandinavian and Northwest European haplotypes. JJ6 is separated by two mutational steps from a modern Finnish haplotype, while JJ1 and JJ2 fall within clusters containing ancient and modern Northwest and Central European sheep.

JJ2 and JJ4 are assigned to haplogroups B1a with ambiguous subclade resolution between B1a1b and B1a2a1. Despite this uncertainty, both samples integrate within the principal B1a component of the median-joining network and do not resolve as geographically isolated.

Closer inspection of the median-joining network (Figure 2) suggests subtle regional patterning; however, this should be interpreted cautiously in light of the overall near-zero population differentiation. Several Danish haplotypes show affinities with the wider Baltic Sea region and Fennoscandian lineages. These affinities reflect individual haplotype relationships within a broadly shared northern European maternal gene pool rather than discrete regional structuring. This includes connections between JJ1 and JJ3 and Finnish haplotypes with a Norwegian lineage positioned basally within that cluster, as well as links between JJ6 and JJ4 and Finnsheep consistent with previously documented connectivity among sheep populations across the Baltic Sea region (Larsson et al., 2024; Rannamäe, Lõugas, Niemi, et al., 2016 b; Rannamäe, Lõugas, Speller, et al., 2016 a).

Although some Danish samples appear in close proximity within the network, these relationships cannot be used to infer intra-site connectivity or geographically structured exchange, given the limited sample size and the constraints of mitochondrial data (Ballard and Whitlock 2004).

None of the Danish haplotypes occupies peripheral or highly divergent positions within the network. This pattern is in alignment with high haplotype diversity (*Hd* = 1.0) and moderate nucleotide diversity (*π* = 0.00123) among the Danish samples, as well as the near-zero differentiation estimates.

## 5 Discussion and conclusions

### 5.1 Maternal lineage diversity in Danish Late Iron Age sheep

The Danish Late Iron Age sheep analysed in this study exhibit high mitochondrial haplotype diversity despite the limited sample size, with each individual representing a distinct maternal lineage within haplogroup B. The absence of haplotype sharing, together with moderate nucleotide diversity and short to moderate mutational distances among sequences, indicates that Danish sheep populations were not maternally homogeneous and instead comprised multiple closely related maternal lineages.

These observations are supported by population differentiation statistics. Pairwise *F*_*ST*_ estimates between Danish ancient sheep and modern Fennoscandian and Northwest European reference populations were low to near-zero, suggesting limited maternal differentiation at the regional scale. Similarly, the median-joining network shows Danish haplotypes dispersed across several branches of the broader European B lineage rather than forming a geographically discrete cluster.

This combination of high haplotype diversity, dispersed network distribution, and low differentiation is not consistent with a recent, severe maternal bottleneck, which would typically reduce mitochondrial diversity through lineage loss (Feinauer et al., 2024). Instead, the data indicate the persistence of multiple maternal lineages within Danish sheep populations during the Germanic Iron Age and Viking Age. Such a pattern is compatible with flock structures in which breeding females were not restricted to a narrow maternal stock and may reflect a combination of local continuity, exchange of animals between communities, and occasional introduction of new breeding females.

Neutrality tests do not reveal significant departures from neutral expectations, although the limited sample size restricts the power of these analyses. Nevertheless, taken together, the diversity statistics, differentiation estimates, and median-joining network topology provide a preliminary genetic baseline for assessing long-term demographic trajectories and regional relationships as additional ancient sheep genomes become available.

Although informative, mitochondrial genomes capture only maternal inheritance and therefore cannot resolve questions regarding nuclear ancestry, effective population size, or male-mediated gene flow (Ballard & Whitlock, 2004). Consequently, the patterns observed here should be interpreted as reflecting maternal lineage diversity rather than a comprehensive demographic history. Moreover, the small sample size limits the detection of subtle demographic events or spatial structuring. Broader temporal sampling and integration with nuclear genomic data will be necessary to evaluate population continuity, replacement, and changes in breeding structure beyond the maternal lineage perspective.

### 5.2 Implications for sheep husbandry and textile production

The maternal lineage diversity observed among Danish Late Iron Age sheep provides a useful genetic perspective on sheep husbandry during a period in which textile production and consumption intensified in Scandinavia. Textile research has demonstrated increased visibility of textile production in the archaeological record during the Late Iron Age, including increasing numbers of textile tools and concentrations of pit houses associated with textile production, as well as an increased demand for textiles related to travel, warfare, and sailing technology (Andersson Strand, 2021). In this context, wool was not simply a household by-product but a key resource embedded within wider economic and social systems of production and exchange. Textile production was organisationally diverse and ranged from household production to household industries, attached specialist production, and workshop-based production (Andersson Strand, 2011).

This is particularly relevant for textiles produced at a large scale, such as sailcloth, where the production demands substantial labour investments and large quantities of textile fibres, such as wool. Estimates of raw material requirements and production time emphasise that textile manufacture requires coordinated systems for raw material procurement, processing, and labour organisation (Andersson Strand, 2016, 2021). Fibre analyses of Viking Age textiles demonstrate substantial variation in wool fibre composition and indicate that textile qualities reflect both the diversity of available fleece and craft-based fibre selection and processing strategies (Andersson Strand, 2011; Andersson Strand & Demant, 2023; Skals et al., 2024).

Within the context of textile production, the mitogenomic data primarily provide a demographic context rather than direct evidence of wool production strategies. The presence of multiple distinct maternal lineages and moderate nucleotide diversity suggests that sheep husbandry systems were not restricted to narrow maternal breeding stocks, a condition that could otherwise have limited flock resilience and productive capacities. Rather, the observed genetic patterns are compatible with flock structures that were not demographically constrained and may have facilitated the maintenance of genetically heterogeneous sheep populations, alongside reproductive stability over time, both of which would have been advantageous in textile economies characterised by increasing scale and organisational complexity. In this context, sheep were likely moved between regions as part of wider systems of exchange, contributing to the maintenance of such diversity. Such exchange may have taken different forms, including both continuous low-level movement and more episodic introductions of animals.

Congruently, mitochondrial genomes capture just maternal inheritance and cannot resolve specific questions regarding livestock mobility, breeding or wool production strategies. The observed patterns of limited maternal differentiation at the broader northern European scale are therefore interpreted as compatible with interactions and exchange between communities rather than as direct evidence of animal movements or structured breeding systems. This interpretation, however, aligns with archaeological models emphasising redistribution, exchange, and regional integration of production at trading sites and central places, where textile production could be embedded within wider economic networks (French et al., 2024; Larsson et al., 2020; Löffelmann et al., 2023; Macheridis et al., 2024).

Taken together, the genetic and archaeological evidence suggests that sheep husbandry during the Late Iron Age was organised within demographic conditions that did not impose strong maternal constraints on flock composition. Rather than indicating narrowly structured breeding populations, the mitogenomic data are consistent with husbandry systems capable of sustaining diverse maternal lineages alongside an expanding textile economy. Mitochondrial variation, therefore, offers insight into maternal population structure relevant to sheep management, while the specific mechanisms underlying breeding strategies and wool production remain better addressed through nuclear genomic data and expanded temporal datasets.

## 6 Data availability

Raw sequence reads are accessible at the Electronic Research Data Archive at the University of Copenhagen (UCPH ERDA). https://erda.ku.dk/archives/112213a5d8a14c3a03f128ee80b6f0fe/published-archive.html

## 7 CRediT authorship contribution statement

**Jonas Holm Jæger:** Conceptualisation, Formal analysis, Investigation, Data curation, Writing - original draft, Writing - review & editing, Visualisation. **Valeria Mattiangeli:** Writing - review & editing, Validation, Investigation. **Jens Ulriksen:** Writing - review & editing, Resources. **Torben Sarauw:** Writing - review & editing, Resources. **Mads Dengsø Jessen:** Writing - review & editing, Resources.

## 8 Declaration of interests

The author declares that they have no known competing financial interests or personal relationships that could have appeared to influence the work reported in this paper.

## 9 Funding

This research is part of the Textile Resources in Viking Age Landscapes project (Centre for Textile Research, University of Copenhagen), funded by the Independent Research Fund Denmark (Grant No. DFF-2027-00204B).

## 10 Acknowledgements

I thank my thesis supervisors, Eva Birgitta Andersson Strand (Centre for Textile Research, University of Copenhagen) and Matthew James Collins (GLOBE Institute, University of Copenhagen), as well as Anne Birgitte Gotfredsen (GLOBE Institute, University of Copenhagen), for their tireless support and invaluable comments on the manuscript. I also want to express my deep gratitude to Daniel Bradley and Aíne Halpin (Smurfit Institute of Genetics, Trinity College Dublin) for providing laboratory facilities and assistance during the laboratory work in Dublin.

I also thank the National Museum of Denmark, Museum Sydøstdanmark, and Nordjyske Museer for granting access to archaeological material used in this study.

## Notes

### Competing Interest Statement

The authors have declared no competing interest.

## Bibliography

Andrews, S. (2010). FastQC: a quality control tool for high throughput sequence data. Available online at: http://www.bioinformatics.babraham.ac.uk/projects/fastqc

Andersson Strand, E. (2011). Tools and Textiles - Production and Organisation in Birka and Hedeby. In S. Sigmundsson (Ed.), Viking Settlements and Viking Society - Papers from the Proceedings of the Sixteenth Viking Congress, Reykjavík and Reykholt, 16th-23rd August 2009. University of Iceland Press.

Andersson Strand, E. (2016). Segel och segelduksproduktion i arkeologisk kontext. In M. Ravn, L. G. Thomsen, E. Andersson Strand, & H. Lyngstrøm (Eds.), Vikingetidens sejl - Festskrift tilegnet Erik Andersen (pp. 21–48). SAXO-Instituttet.

Andersson Strand, E. (2021). Weaving textiles: Textile consumption for travel and warfare. Viking, 84(1), 167–186. 10.5617/viking.9053

Andersson Strand, E., & Demant, I. (2023). Fibres, Tools & Textiles. Nationalmuseet.

Ballard, J. W. O., & Whitlock, M. C. (2004). The incomplete natural history of mitochondria. Molecular Ecology, 13(4), 729–744. 10.1046/j.1365-294x.2003.02063.x

Bandelt, H. J., Forster, P., & Röhl, A. (1999). Median-joining networks for inferring intraspecific phylogenies. Molecular Biology and Evolution, 16(1), 37–48. 10.1093/oxfordjournals.molbev.a026036

Boessenkool, S., Hanghøj, K., Nistelberger, H. M., Der Sarkissian, C., Gondek, A. T., Orlando, L., Barrett, J. H., & Star, B. (2016). Combining bleach and mild predigestion improves ancient DNA recovery from bones. Molecular Ecology Resources, 17(4), 742–751. 10.1111/1755-0998.12623

Brandt, L. Ø. (2014). Species identification of skins and development of sheep wool: An interdisciplinary study combining textile research, archaeology, and biomolecular methods [Ph.D. thesis]. University of Copenhagen.

Chessa, B., Pereira, F., Arnaud, F., Amorim, A., Goyache, F., Mainland, I., Kao, R. R., Pemberton, J. M., Beraldi, D., Stear, M. J., Alberti, A., Pittau, M., Iannuzzi, L., Banabazi, M. H., Kazwala, R. R., Zhang, Y.-P., Arranz, J. J., Ali, B. A., Wang, Z., … Palmarini, M. (2009). Revealing the History of Sheep Domestication Using Retrovirus Integrations. Science, 324(5926), 532–536. 10.1126/science.1170587

Dabney, J., Knapp, M., Glocke, I., Gansauge, M.-T., Weihmann, A., Nickel, B., Valdiosera, C., García, N., Pääbo, S., Arsuaga, J.-L., & Meyer, M. (2013). Complete mitochondrial genome sequence of a Middle Pleistocene cave bear reconstructed from ultrashort DNA fragments. Proceedings of the National Academy of Sciences of the United States of America, 110(39), 15758–15763. 10.1073/pnas.1314445110

Daly, K. G., Mullin, V. E., Hare, A. J., Halpin, Á., Mattiangeli, V., Teasdale, M. D., Rossi, C., Geiger, S., Krebs, S., Medugorac, I., Sandoval-Castellanos, E., Özbaşaran, M., Duru, G., Gülcür, S., Pöllath, N., Collins, M., Frantz, L., Vila, E., Zidarov, P., … Bradley, D. G. (2025). Ancient genomics and the origin, dispersal, and development of domestic sheep. Science, 387(6733), 492–497. 10.1126/science.adn2094

Danecek, P., Bonfield, J. K., Liddle, J., Marshall, J., Ohan, V., Pollard, M. O., Whitwham, A., Keane, T., McCarthy, S. A., Davies, R. M., & Li, H. (2021). Twelve years of SAMtools and BCFtools. GigaScience, 10(2), 1–4. 10.1093/gigascience/giab008

Deng, J., Xie, X.-L., Wang, D.-F., Zhao, C., Lv, F.-H., Li, X., Yang, J., Yu, J.-L., Shen, M., Gao, L., Yang, J.-Q., Liu, M.-J., Li, W.-R., Wang, Y.-T., Wang, F., Li, J.-Q., Hehua, E., Liu, Y.-G., Shen, Z.-Q., … Li, M.-H. (2020). Paternal Origins and Migratory Episodes of Domestic Sheep. Current Biology: CB, 30(20), 4085–4095.e6. 10.1016/j.cub.2020.07.077

Enghoff, I. B. (2018). 6.c.13. Fauna. In J. Ulriksen (Ed.), Vester Egesborg - En anløbsog togtsamlingsplads fra yngre germansk jernalder og vikingetid på Sydsjælland (pp. 286–295). Aarhus Universitetsforlag.

Feinauer, I. S., Lord, E., von Seth, J., Xenikoudakis, G., Ersmark, E., Dalén, L., & Meleg, I.-N. (2024). Heterochronous mitogenomes shed light on the Holocene history of the Scandinavian brown bear. Scientific Reports, 14(1), 1–11. 10.1038/s41598-024-75028-6

French, K. M., Musiał, A. D., Karczewski, M., Daugnora, L., Shiroukhov, R., Ropka-Molik, K., Baranowski, T., Bertašius, M., Skvortsov, K., Szymański, P., Mellin-Wyczółkowska, I., Gręzak, A., Wyczółkowski, D., Pluskowski, A., Andersen, M., Millet, M.-A., Inglis, E., & Madgwick, R. (2024). Biomolecular evidence reveals mares and long-distance imported horses sacrificed by the last pagans in temperate Europe. Science Advances, 10(20). 10.1126/sciadv.ado3529

Gamba, C., Jones, E. R., Teasdale, M. D., McLaughlin, R. L., Gonzalez-Fortes, G., Mattiangeli, V., Domboróczki, L., Kővári, I., Pap, I., Anders, A., Whittle, A., Dani, J., Raczky, P., Higham, T. F. G., Hofreiter, M., Bradley, D. G., & Pinhasi, R. (2014). Genome flux and stasis in a five millennium transect of European prehistory. Nature Communications, 5, 5257. 10.1038/ncomms6257

Groß, D., Presslee, S., Schmölcke, U., Nikulina, E. A., & Hendy, J. (2024). Denmark’s Not-So-Oldest Sheep: An Update on Domestic Animals from the Femern Project. Danish Journal of Archaeology, 13(1), 1–10. 10.7146/dja.v13i1.145009

Hiendleder, S., Mainz, K., Plante, Y., & Lewalski, H. (1998). Analysis of mitochondrial DNA indicates that domestic sheep are derived from two different ancestral maternal sources: no evidence for contributions from urial and argali sheep. The Journal of Heredity, 89(2), 113–120. 10.1093/jhered/89.2.113

Hudson, R. R., Slatkin, M., & Maddison, W. P. (1992). Estimation of levels of gene flow from DNA sequence data. Genetics, 132(2), 583–589. 10.1093/genetics/132.2.583

Jónsson, H., Ginolhac, A., Schubert, M., Johnson, P. L. F., & Orlando, L. (2013). mapDamage2.0: fast approximate Bayesian estimates of ancient DNA damage parameters. Bioinformatics (Oxford, England), 29(13), 1682–1684. 10.1093/bioinformatics/btt193

Jørgensen, L., Thomsen, L. G., & Jørgensen, A. N. (2019). Accommodating assemblies, as evidenced at the 6th–11th-century AD royal residence at lake Tissø, Denmark. In Power and Place in Europe in the Early Middle Ages (pp. 148–173). British Academy. 10.5871/bacad/9780197266588.003.0007

Kaptan, D., Atağ, G., Vural, K. B., Morell Miranda, P., Akbaba, A., Yüncü, E., Buluktaev, A., Abazari, M. F., Yorulmaz, S., Kazancı, D. D., Küçükakdağ Doğu, A., Çakan, Y. G., Özbal, R., Gerritsen, F., De Cupere, B., Duru, R., Umurtak, G., Arbuckle, B. S., Baird, D., … Özer, F. (2024). The population history of domestic sheep revealed by paleogenomes. Molecular Biology and Evolution, 41(10), 1–20. 10.1093/molbev/msae158

Katoh, K., & Standley, D. M. (2013). MAFFT multiple sequence alignment software version 7: improvements in performance and usability. Molecular Biology and Evolution, 30(4), 772–780. 10.1093/molbev/mst010

Korneliussen, T. S., Albrechtsen, A., & Nielsen, R. (2014). ANGSD: Analysis of next generation sequencing data. BMC Bioinformatics, 15(1), 356. 10.1186/s12859-014-0356-4

Kveiborg, J. (2019). Husdyrhold og fiskeri i yngre jernalder. En zooarkæologisk analyse af knoglemateriale fra Bejsebakken. In T. Sarauw (Ed.), Bejsebakken - En nordjysk bebyggelse fra yngre jernalder og vikingetid (Vol. 29, pp. 199–215).

Larsson, M., Magnell, O., Styring, A., Lagerås, P., & Evans, J. (2020). Movement of agricultural products in the Scandinavian Iron Age during the first millennium AD: 87Sr/86Sr values of archaeological crops and animals in southern Sweden. Science and Technology of Archaeological Research, 6(1), 96–112. 10.1080/20548923.2020.1840121

Larsson, M. N. A., Miranda, P. M., Pan, L., Vural, K. B., Kaptan, D., Soares, A. E. R., Kivikero, H., Kantanen, J., Somel, M., Özer, F., Johansson, A. M., Storå, J., & Günther, T. (2024). Ancient sheep genomes reveal four Millennia of north European short-tailed sheep in the Baltic Sea region. Genome Biology and Evolution, 16(6), evae114. 10.1093/gbe/evae114

Leigh, J. W., & Bryant, D. (2015). Popart: Full-feature software for haplotype network construction. Methods in Ecology and Evolution, 6(9), 1110–1116. 10.1111/2041-210x.12410

Li, H., & Durbin, R. (2009). Fast and accurate short read alignment with Burrows-Wheeler transform. Bioinformatics (Oxford, England), 25(14), 1754–1760. 10.1093/bioinformatics/btp324

Löffelmann, T., Snoeck, C., Richards, J. D., Johnson, L. J., Claeys, P., & Montgomery, J. (2023). Sr analyses from only known Scandinavian cremation cemetery in Britain illuminate early Viking journey with horse and dog across the North Sea. PloS One, 18(2), e0280589. 10.1371/journal.pone.0280589

Macheridis, S., Faillace, K., Hood, M., Sayle, K. L., Inglis, E., & Madgwick, R. (2024). Sheep Ahoy: Exploring sheep management and its role in Viking Age economy through multiproxy analyses at Löddeköpinge, Sweden. International Journal of Osteoarchaeology. 10.1002/oa.3355

MacHugh, D. E., Edwards, C. J., Bailey, J. F., Bancroft, D. R., & Bradley, D. G. (2000). The Extraction and Analysis of Ancient DNA From Bone and Teeth: a Survey of Current Methodologies. Ancient Biomolecules, 3, 81–102.

Meadows, J. R. S., Cemal, I., Karaca, O., Gootwine, E., & Kijas, J. W. (2007). Five Ovine Mitochondrial Lineages Identified From Sheep Breeds of the Near East. Genetics, 175(3), 1371–1379. 10.1534/genetics.106.068353

Meyer, M., & Kircher, M. (2010). Illumina sequencing library preparation for highly multiplexed target capture and sequencing. Cold Spring Harbor Protocols, 2010(6), db.prot5448. 10.1101/pdb.prot5448

Morell Miranda, P., Soares, A. E. R., & Günther, T. (2023). Demographic reconstruction of the Western sheep expansion from whole-genome sequences. G3 (Bethesda, Md.), 13(11). 10.1093/g3journal/jkad199

Pääbo, S., Poinar, H., Serre, D., Jaenicke-Despres, V., Hebler, J., Rohland, N., Kuch, M., Krause, J., Vigilant, L., & Hofreiter, M. (2004). Genetic analyses from ancient DNA. Annual Review of Genetics, 38(1), 645–679. 10.1146/annurev.genet.37.110801.143214

Peng, M.-S., Fan, L., Shi, N.-N., Ning, T., Yao, Y.-G., Murphy, R. W., Wang, W.-Z., & Zhang, Y.-P. (2015). DomeTree: a canonical toolkit for mitochondrial DNA analyses in domesticated animals. Molecular Ecology Resources, 15(5), 1238–1242. 10.1111/1755-0998.12386

Rannamäe, E., Lõugas, L., Niemi, M., Kantanen, J., Maldre, L., Kadõrova, N., & Saarma, U. (2016). Maternal and paternal genetic diversity of ancient sheep in Estonia from the Late Bronze Age to the post-medieval period and comparison with other regions in Eurasia. Animal Genetics, 47(2), 208–218. 10.1111/age.12407

Rannamäe, E., Lõugas, L., Speller, C. F., Valk, H., Maldre, L., Wilczyński, J., Mikhailov, A., & Saarma, U. (2016). Three Thousand Years of Continuity in the Maternal Lineages of Ancient Sheep (Ovis aries) in Estonia. PloS One, 11(10), e0163676. 10.1371/journal.pone.0163676

Rannamäe, E., Saarma, U., Ärmpalu-Idvand, A., Teasdale, M. D., & Speller, C. (2020). Retroviral analysis reveals the ancient origin of Kihnu native sheep in Estonia: implications for breed conservation. Scientific Reports, 10(1), 17340. 10.1038/s41598-020-74415-z

Roesdahl, E., Sindbæk, S. M., & Pedersen, A. (Eds.). (2014). Aggersborg i vikingetiden: bebyggelsen og borgen. Aarhus University Press.

Rosenlund, K., & Hatting, T. (2014). Zoological finds. In E. Roedahl, S. M. Sindbæk, A. Pedersen, & D. M. Wilson (Eds.), Aggersborg: The Viking Age Settlement and Fortress (pp. 373–381). Jutland Archaeological Soceity Publications.

Rozas, J., Ferrer-Mata, A., Sánchez-DelBarrio, J. C., Guirao-Rico, S., Librado, P., Ramos-Onsins, S. E., & Sánchez-Gracia, A. (2017). DnaSP 6: DNA Sequence Polymorphism analysis of large data sets. Molecular Biology and Evolution, 34(12), 3299–3302. 10.1093/molbev/msx248

Sarauw, T. (2019). Bejsebakken: en nordjysk bebyggelse fra yngre jernalder og vikingetid. University Press of Southern Denmark.

Schroeder, O., Benecke, N., Frölich, K., Peng, Z., Kaniuth, K., Sverchkov, L., Reinhold, S., Belinskiy, A., & Ludwig, A. (2017). Endogenous Retroviral Insertions Indicate a Secondary Introduction of Domestic Sheep Lineages to the Caucasus and Central Asia between the Bronze and Iron Age. Genes, 8(6). 10.3390/genes8060165

Schubert, M., Lindgreen, S., & Orlando, L. (2016). AdapterRemoval v2: rapid adapter trimming, identification, and read merging. BMC Research Notes, 9(1), 88. 10.1186/s13104-016-1900-2

Skals, I., Rimstad, C., & Mannering, U. (2024). Viking Age Wool Fibres. Nationalmuseet.

Skoglund, P., Ersmark, E., Palkopoulou, E., & Dalén, L. (2015). Ancient wolf genome reveals an early divergence of domestic dog ancestors and admixture into high-latitude breeds. Current Biology: CB, 25(11), 1515–1519. 10.1016/j.cub.2015.04.019

Sørensen, L. (2014). From Hunter to Farmer in Northern Europe - Migration and Adaptation during the Neolithic and Bronze Age. Acta Archaeologica, 85(1). 10.1111/j.1600-0390.2014.00945.x

Ulriksen, J. (2018). Vester Egesborg - En anløbsog togtsamlingsplads fra yngre germansk jernalder og vikingetid på Sydsjælland (Vol. 1). Aarhus Universitetsforlag. 10.2307/j.ctv34wmrfz

Vigne, J.-D. (2011). The origins of animal domestication and husbandry: a major change in the history of humanity and the biosphere. Comptes Rendus Biologies, 334(3), 171–181. 10.1016/j.crvi.2010.12.009

Wingett, S. W., & Andrews, S. (2018). FastQ Screen: A tool for multi-genome mapping and quality control. F1000Research, 7, 1338. 10.12688/f1000research.15931.2

Yang, D. Y., Eng, B., Waye, J. S., Dudar, J. C., & Saunders, S. R. (1998). Technical note: improved DNA extraction from ancient bones using silica-based spin columns. American Journal of Physical Anthropology, 105(4), 539–543.

Yurtman, E., Özer, O., Yüncü, E., Dağtaş, N. D., Koptekin, D., Çakan, Y. G., Özkan, M., Akbaba, A., Kaptan, D., Atağ, G., Vural, K. B., Gündem, C. Y., Martin, L., Kılınç, G. M., Ghalichi, A., Açan, S. C., Yaka, R., Sağlıcan, E., Lagerholm, V. K., … Özer, F. (2021). Archaeogenetic analysis of Neolithic sheep from Anatolia suggests a complex demographic history since domestication. Communications Biology, 4(1), 1279. 10.1038/s42003-021-02794-8

Zeder, M. A. (2008). Domestication and early agriculture in the Mediterranean Basin: Origins, diffusion, and impact. Proceedings of the National Academy of Sciences of the United States of America, 105(33), 11597–11604. 10.1073/pnas.0801317105

