## Supplementary S2 for "Mitochondrial genomes reveal maternal lineages of Late Iron Age sheep (*Ovis aries*) in Denmark"

### **S2. Diagnostic coverage plots used for molecular sex estimation of ancient individuals.**

**JJ1**
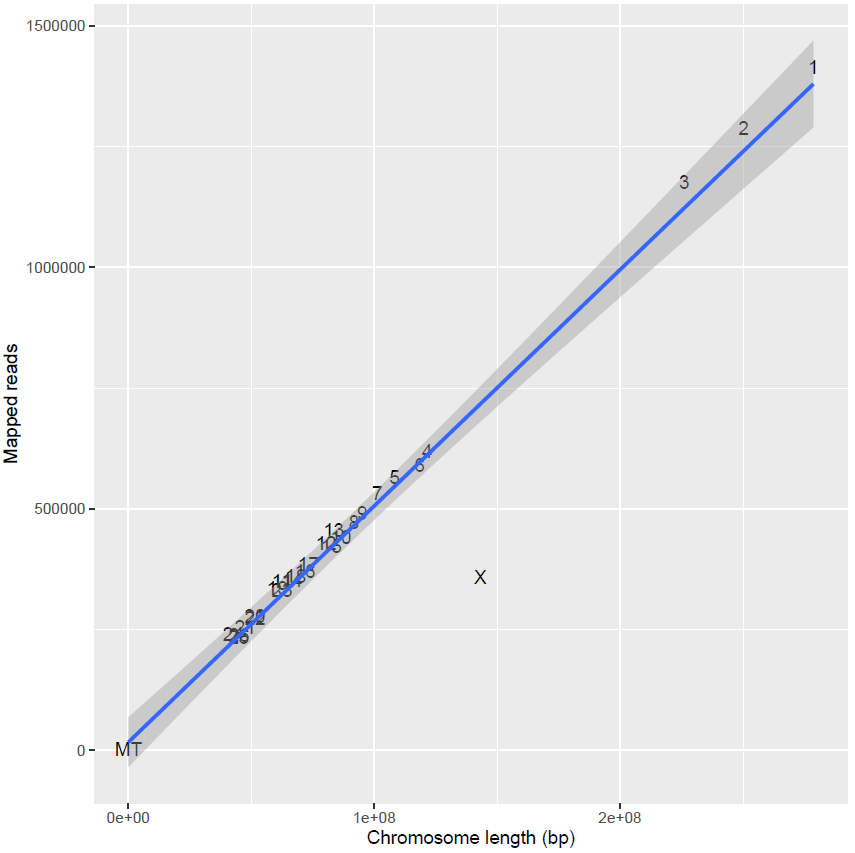

**JJ2**
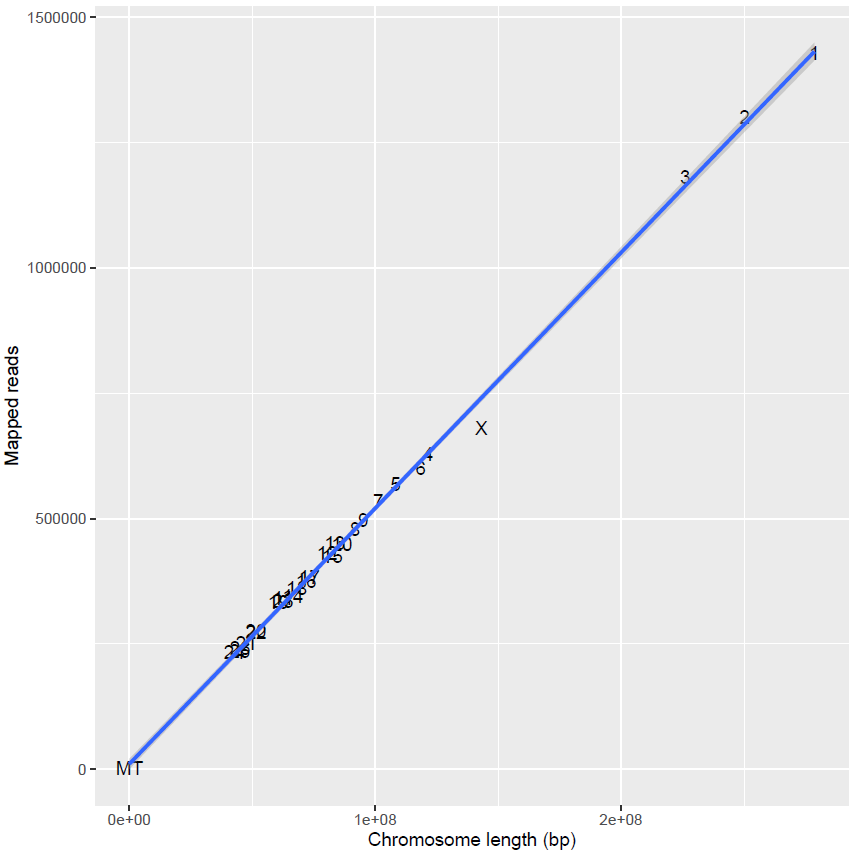

**JJ3**
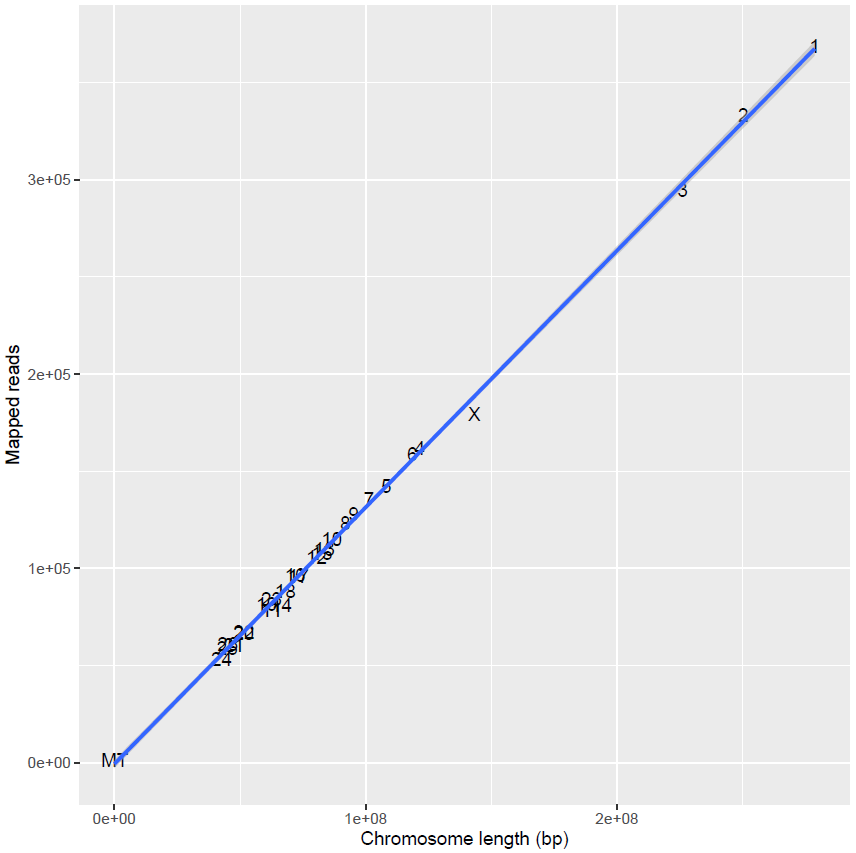

**JJ4**
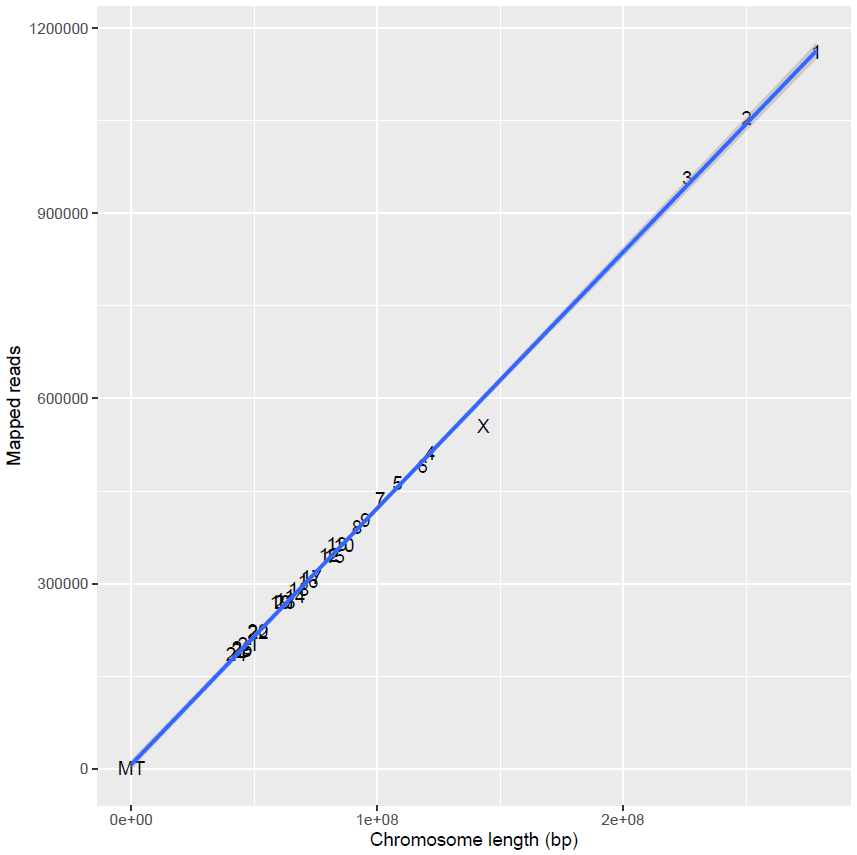

**JJ5**
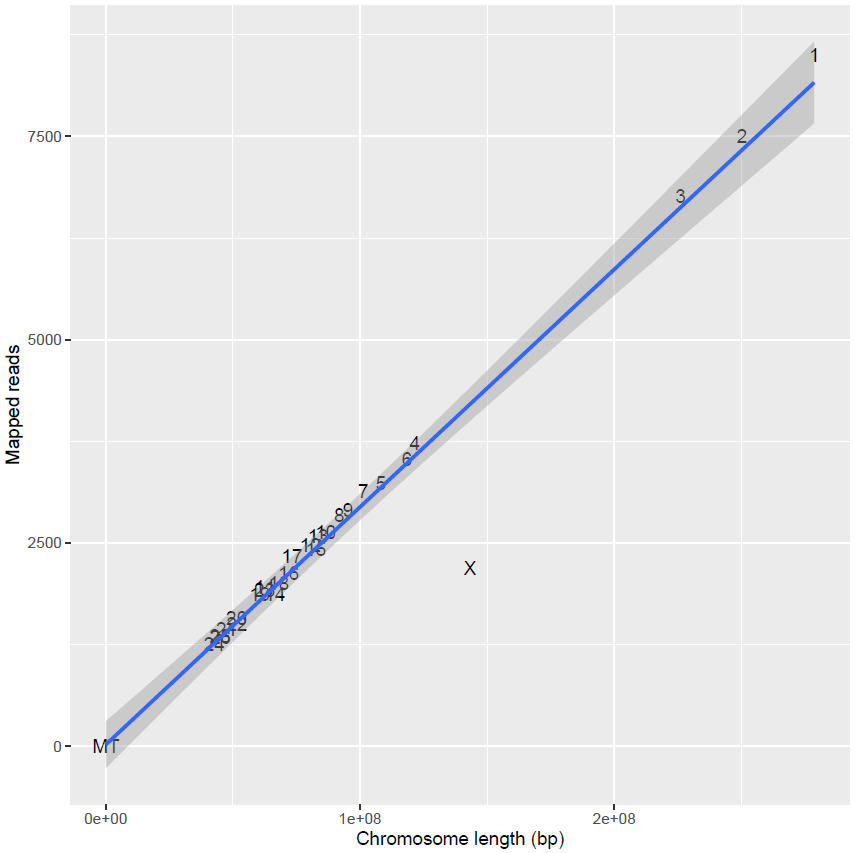

**JJ6**
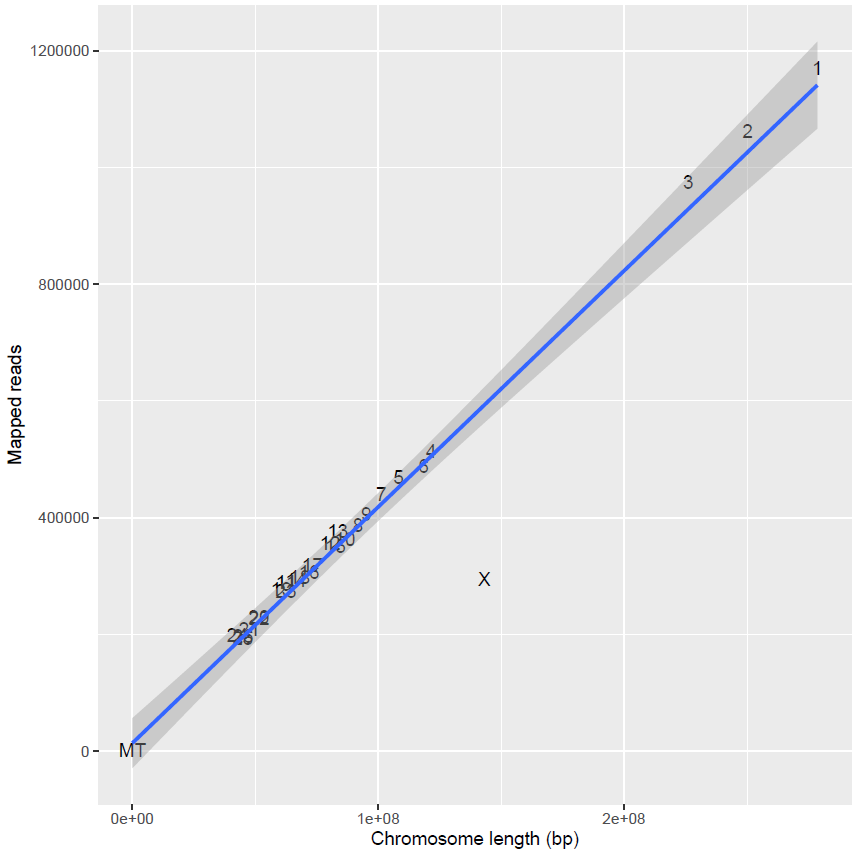
