## Supplementary figures and images for "Mitochondrial genomes reveal maternal lineages of Late Iron Age sheep (*Ovis aries*) in Denmark"

### Supplementary S4

#### **S4. Ancient DNA damage and length distribution plots for all newly generated ancient samples.**

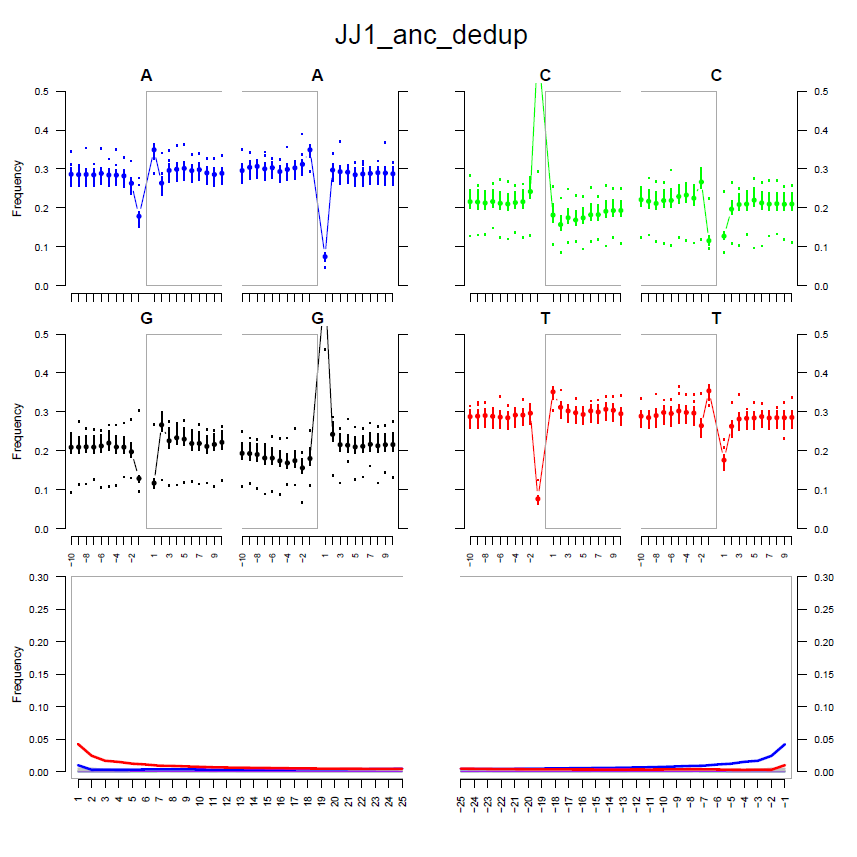

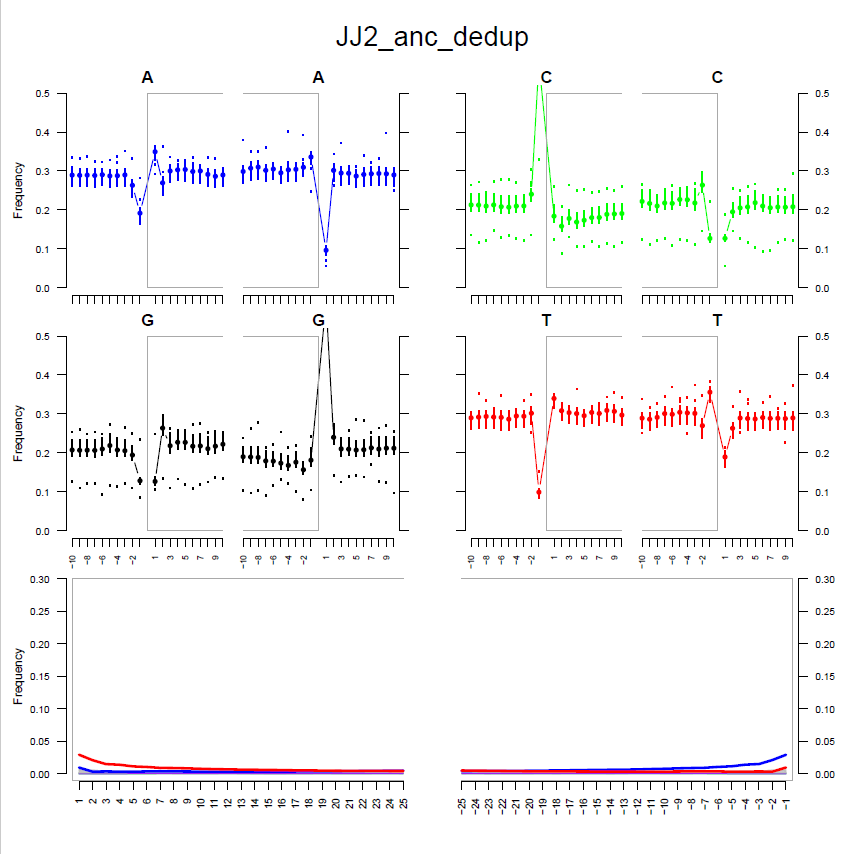

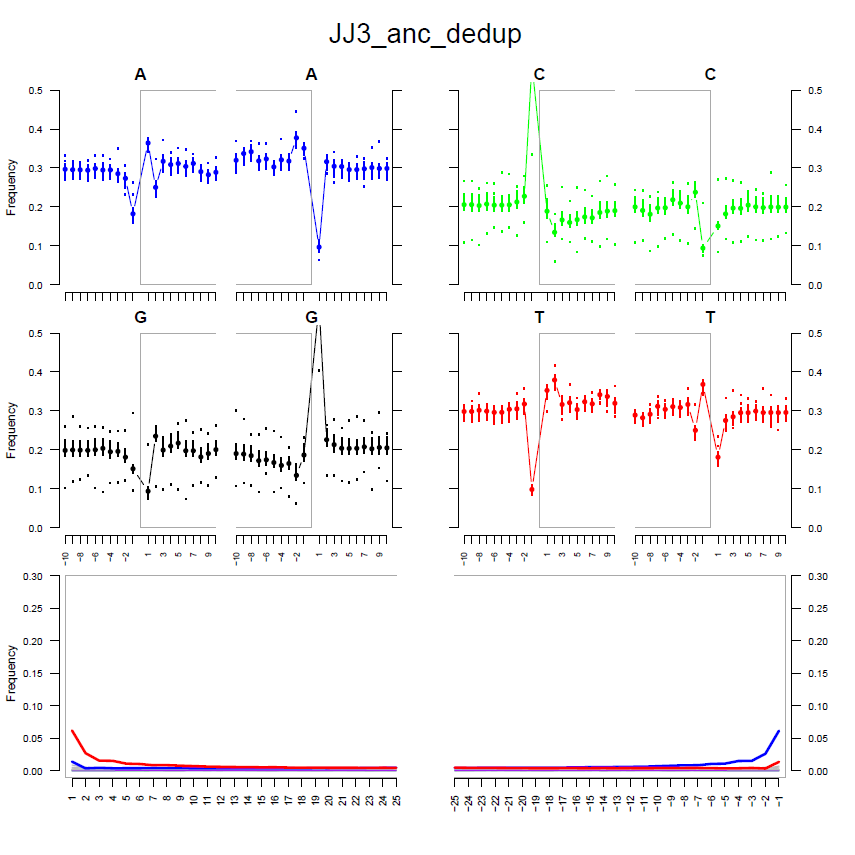

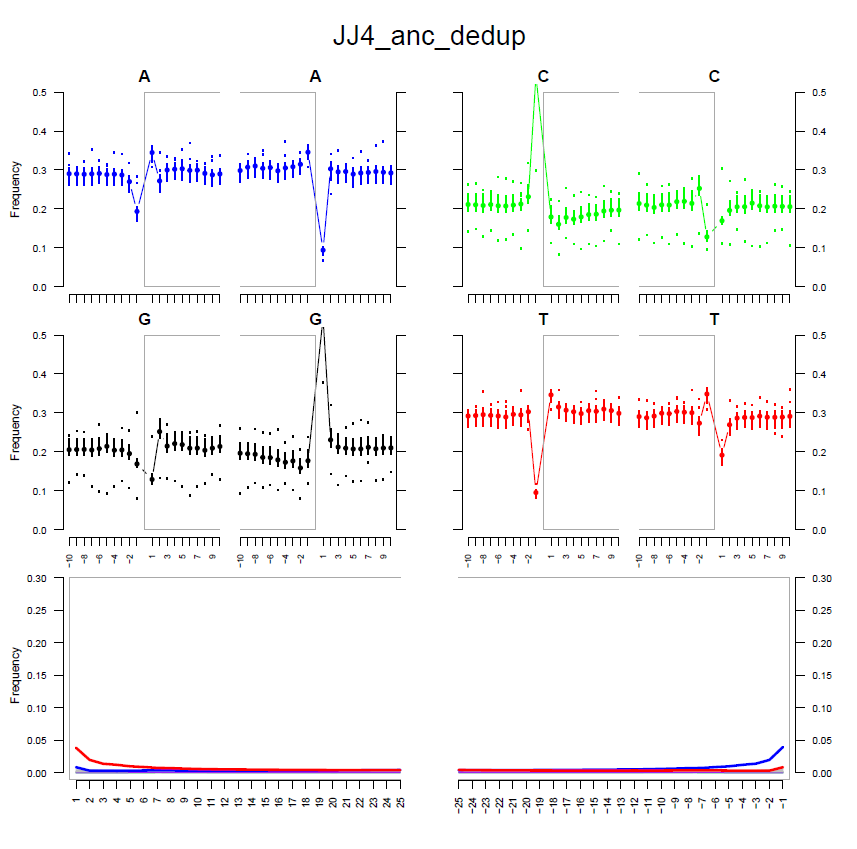

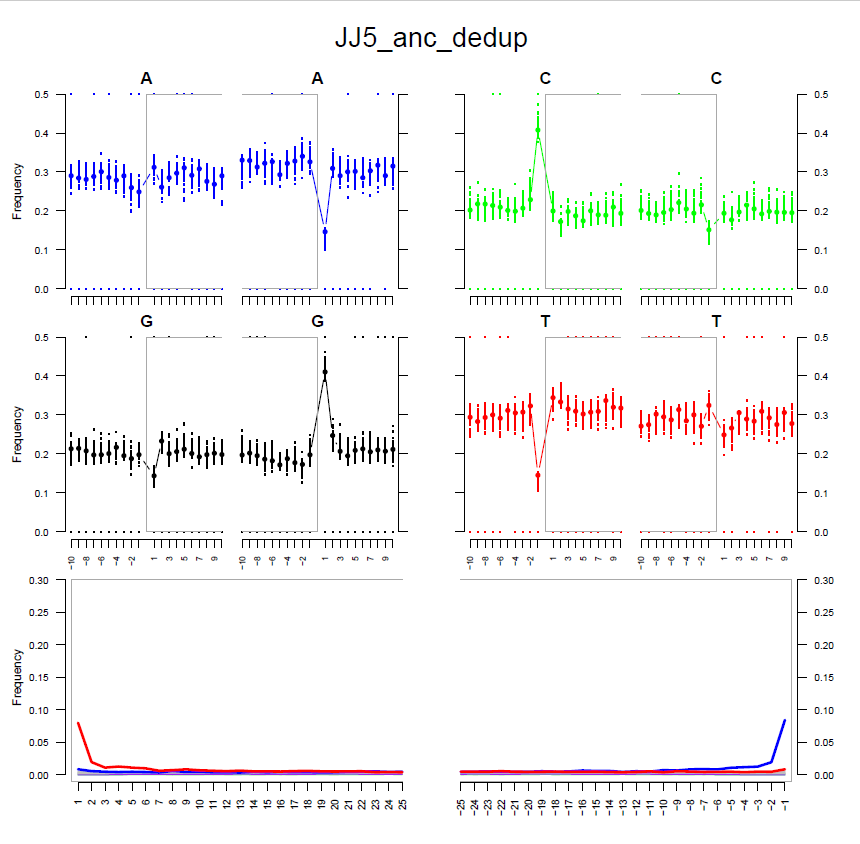

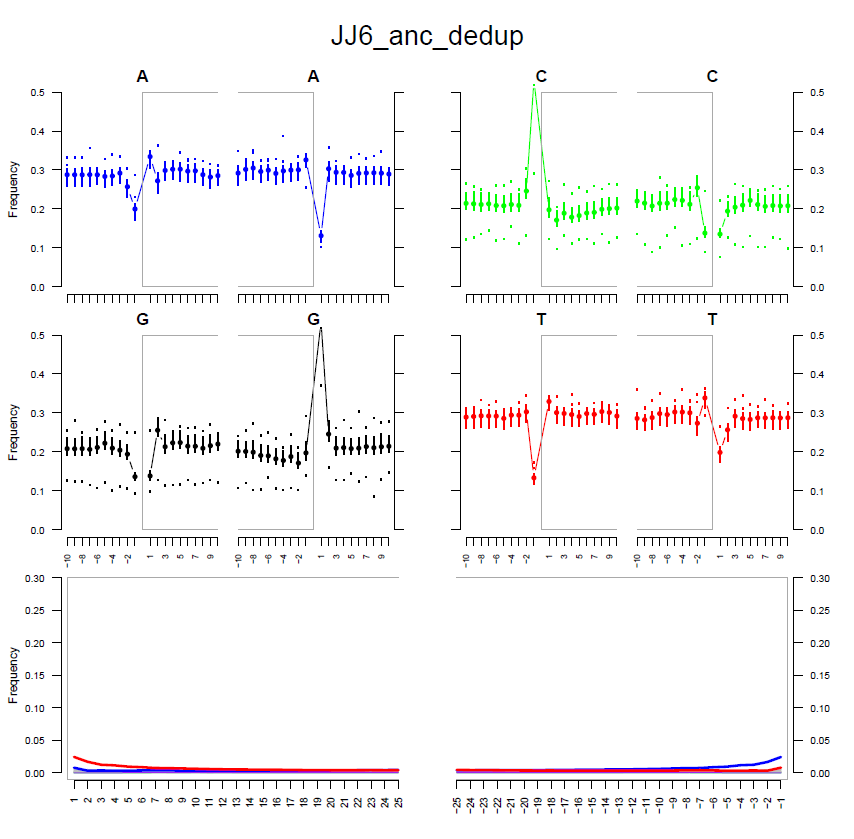

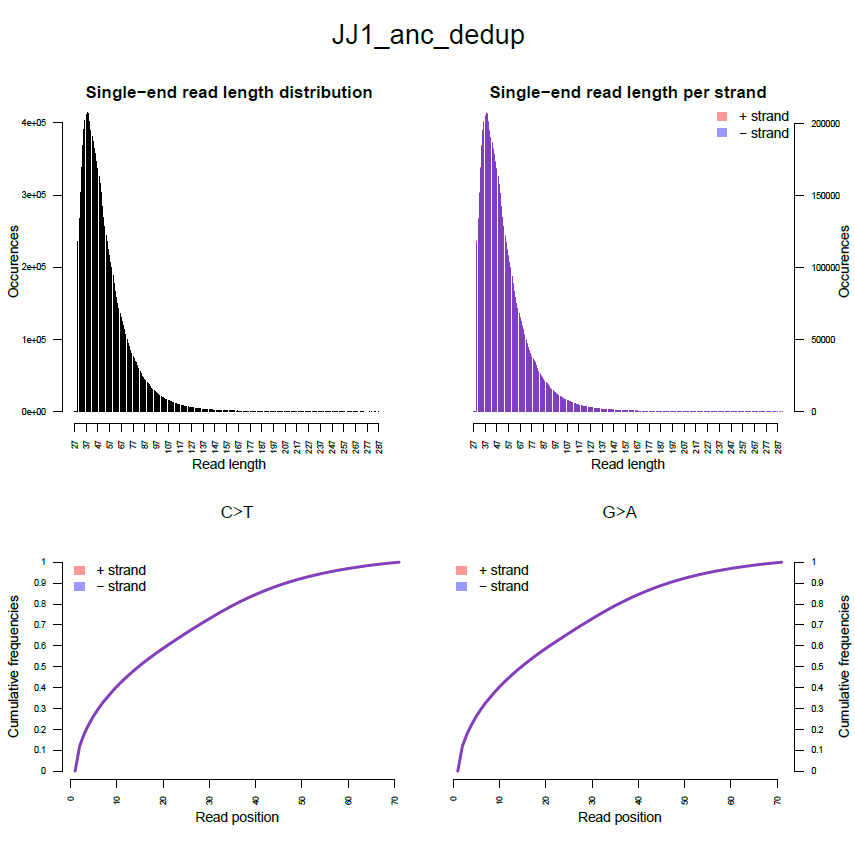

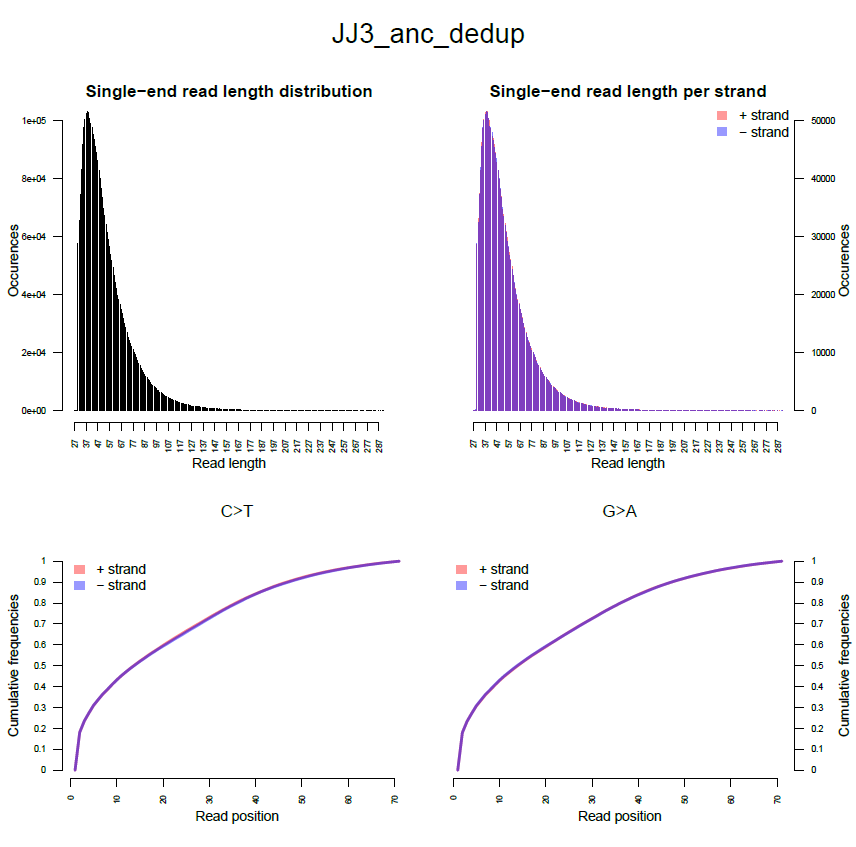

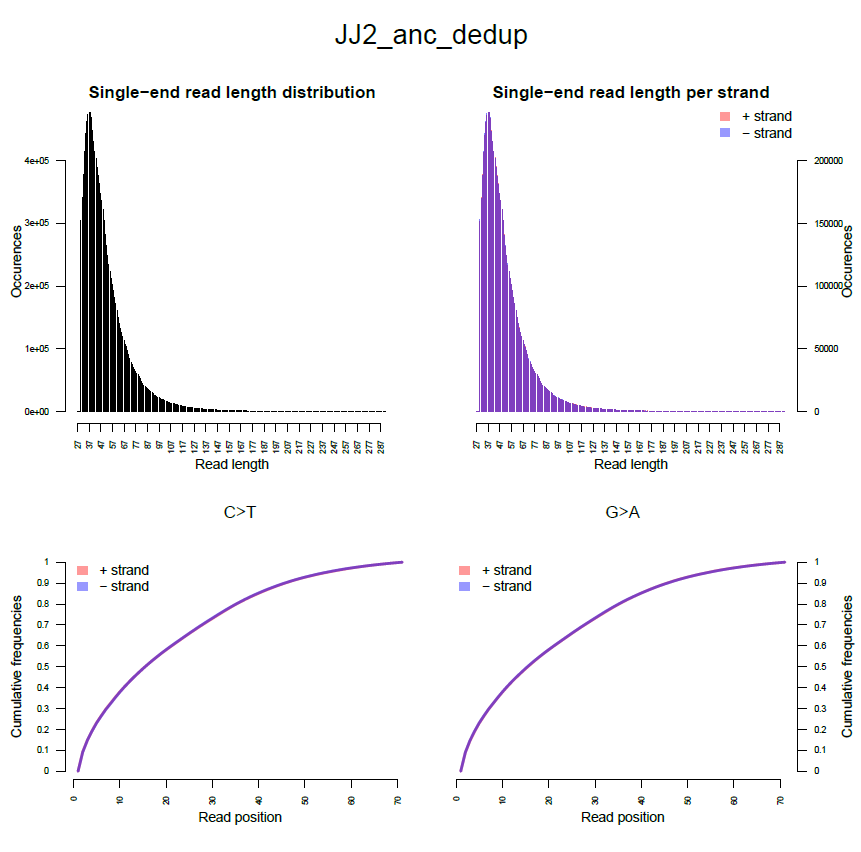

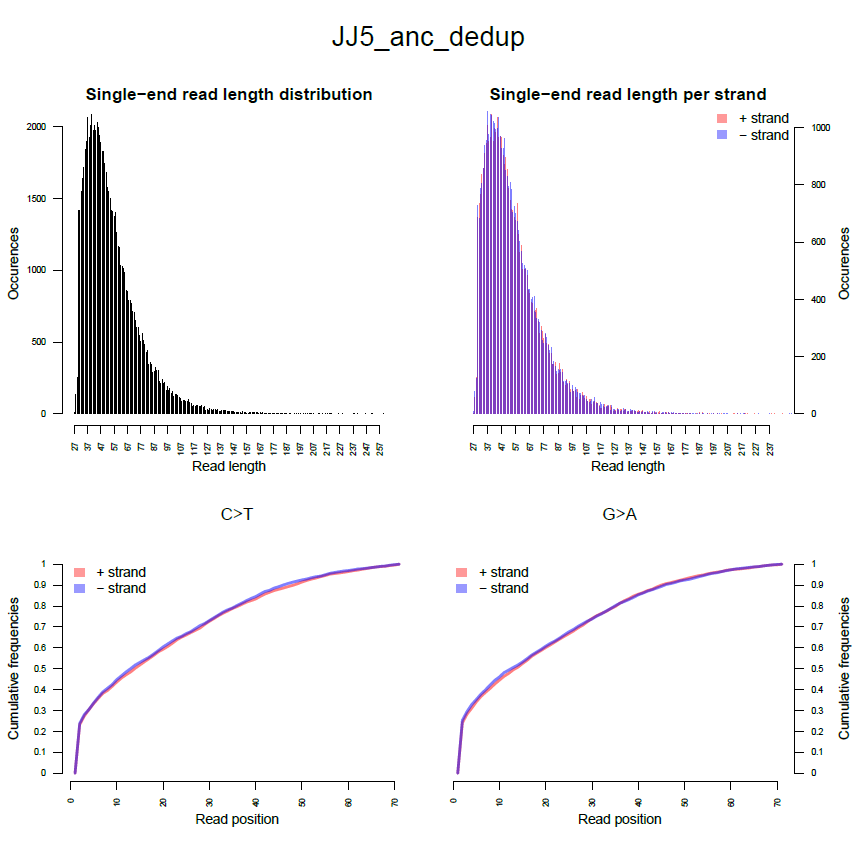

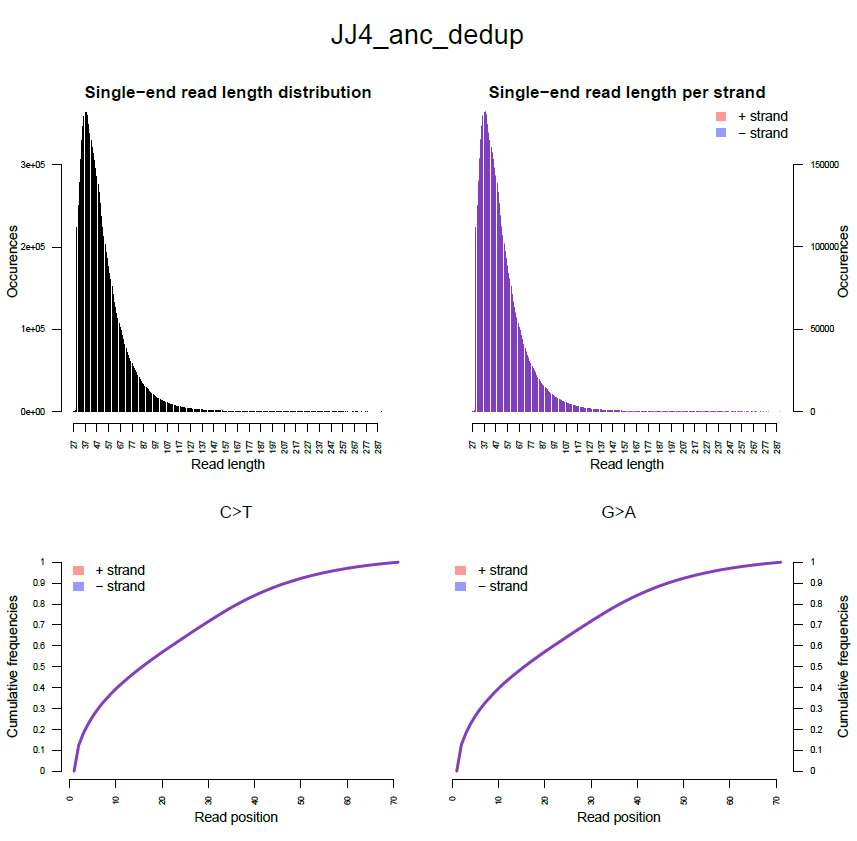

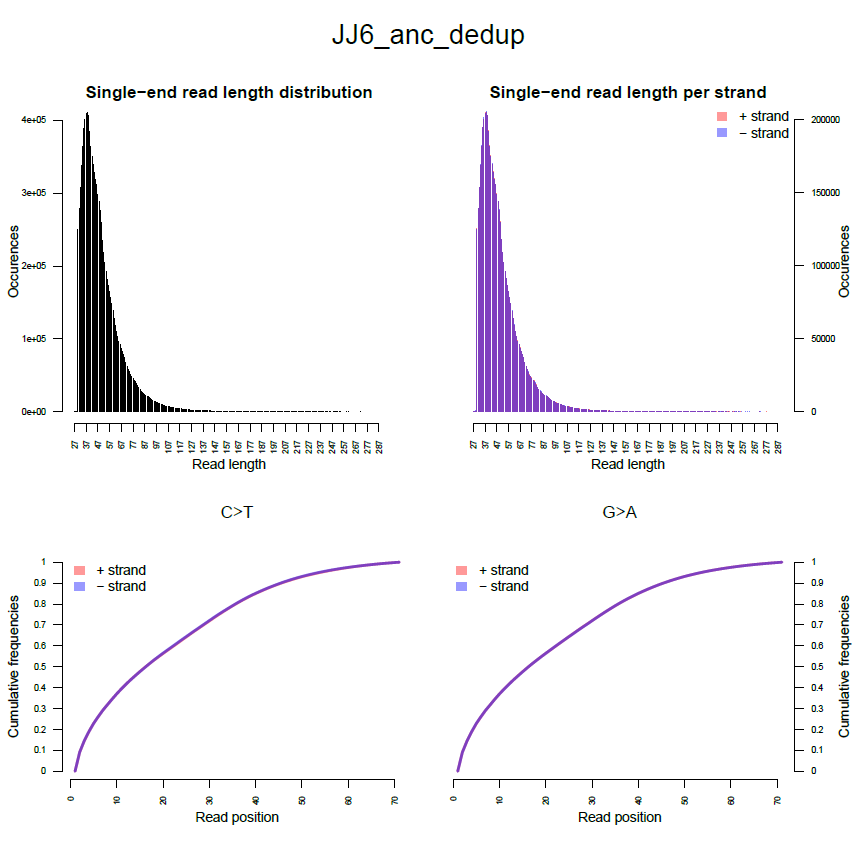
